# Effects of Chinese herbal medicine residues on antibiotic resistance genes and the bacterial community in chicken manure composting

**DOI:** 10.1101/796086

**Authors:** Jinping Wu, Junjian Li, Jianwen Chen, Dale Li, Hong Zhang, Zhenyu Li

## Abstract

Livestock manure is an important way that antibiotic resistance genes (ARGs) can enter the environment, and composting is an effective method for removing ARGs from livestock manure. In this study, different volume ratios of Chinese medicinal herbal residues (CMHRs) were added to laboratory-scale chicken manure composting to evaluate their effects on the behavior of ARGs, mobile genetic elements (MGEs), and the bacterial community. At the end of the composting period, the structure of the microbial community changed. Firmicutes decreased and Bacteroidetes increased. The relative abundance of the 21 ARGs and 5 MGEs detected decreased by varying degrees in the different treatments (except for *sul*I and *int*I1). The removal rate of the ARGs increased with the increased addition of CMHRs. The correlations between transferase genes (*tnp*A and *tnp*A-02) and ARGs were significant (*p* < 0.05); therefore, transposon plays an important role in the horizontal gene transfer of ARGs in chicken manure. The results imply that CMHRs would be an effective bulking agent for the removal of ARGs from chicken manure composting.

## Introduction

Antibiotics are widely used in livestock cultivation, with 52% of antibiotics are used as growth promoters and agents (Zhang et al., 2015; Zhao et al., 2018). Veterinary antibiotics used during livestock cultivation are a major source of antibiotic pollution in the soil environment; the main way they enter the soil is by farmers directly applying feces or urine containing undecomposed veterinary drugs or metabolic intermediates of veterinary drugs onto the soil as organic fertilizers (Heuer et al., 2011; Han et al., 2018). The long-term and excessive non-standard application of antibiotics to the environment leads to the presence of an increased amount of residual antibiotics, thus increasing bacterial resistance and drug-resistant strains in the environment. The spread of bacterial resistance is closely related to an increase in the presence of environmental resistant microorganisms (Martinez et al., 2009). Bacterial resistance is mainly manifested by the expression of antibiotic resistance genes (ARGs) in bacteria, which are transmitted between different species and environments via horizontal gene transfer through mobile genetic elements (MGEs), such as plasmids, transposons, and integrons (Wright, 2007; Zhang et al., 2016; Zhao et al., 2018). In addition, previous studies confirmed that livestock manure facilitates the horizontal transfer of ARGs in microbes in the soil. ARGs in livestock manure can be further transferred from the environment to human beings through the food chain, thereby increasing the risk of obtaining bacterial infectious diseases and seriously threatening human health (Zhou et al., 2007).

According to 2005 estimates, China is the world’s largest livestock and poultry breeding country, with approximately 19 billion tons of livestock manure being produced annually (Qiu et al., 2013). More than 80% of these livestock manure without integrated treatment was directly applied to farmland, therefore, there is a high risk to ecological and environmental security (Laguë et al., 2005). To prevent the accumulation of ARGs in livestock manure for agricultural use, the concentration of ARGs in the manure should be decreased to mitigate environmental risks. Recent studies shown that composting can be used to decrease the abundance of ARGs in livestock manure (Chen et al., 2007; Duan et al., 2018). Composting also changed the microbial community structure that affects the abundance of ARGs. Studies shown that many quinolone resistance genes (Cui et al., 2016), tetracycline resistance genes (Ma et al., 2011), and some beta-lactam resistance genes (Zhang et al., 2016) were reduced after composting, and the bacterial community significantly changed. The different fates of ARGs in livestock manure may be associated with the composition of the composting bulking agent or the bacterial structure during the composting process (Tien et al., 2017).

It is a common composting method to add a bulking agent to livestock manure compost, which adjusts the humidity of the livestock manure and the ratio of carbon and nitrogen to reach ideal composting conditions, and rapidly increases the composting temperature. The Chinese herbal medicine industry is developing rapidly, and thus many Chinese medicinal herbal residues (CMHRs) have been produced. The annual output of CMHRs has reached up to 30 million tons, and it is still on the rise in China (Zhang et al., 2016). Chang et al. (2010) found that CMHRs improved soil physical and chemical properties, and CMHRs with appropriate amounts of inorganic fertilizer promoted corn root growth. Wang et al. (2008) found that aerobic thermophilic compost technology is a feasible way to treat solid slag of CMHRs.

Most of the research on the changes of ARGs during the composting process has focused on the use of bamboo charcoal, sawdust, rice husk, wheat, cotton stalks, or biochars as the compost-bulking agents (Duan et al., 2018; Li et al., 2017; Zhang et al., 2019), however, CMHRs were rarely used. Chicken manure occupies a large proportion in livestock manure, and the relative abundance of ARGs in chicken manure is higher than that in swine manure (Han et al., 2018). In the this study, we aimed to explore the effects of composting on the microbial community and abundance of ARGs by the addition of different amounts of CMHRs in chicken manure, and to select the best composting mode for effectively decreasing the abundance of ARGs.

## Materials and methods

### Experimental design

The chicken manure used in this experiment was obtained from a livestock farm and the CMHRs were collected from a pharmaceutical factory in Taiyuan, China. The composting experiment was conducted in an outdoor composting bin. Different doses of CMHRs were added to chicken manure for composting, with four composting treatments established with different volume ratios as follows: (1) CK: only chicken manure without the addition of any bulking agent, (2) M1: chicken manure with 50% (v/v) CMHRs added, (3) M2: chicken manure with 30% (v/v) CMHRs added, and (4) M3: chicken manure with 20% (v/v) CMHRs added. The temperature was measured using a thermometer at the center of each compost mixture at 8:00 each day, and the environmental temperature was recorded at the same time.

### Sample collection

Samples were collected by mixing the top, middle, and bottom samples of the compost and rolling the container to achieve uniform distribution. The samples collected on days 0, 3, 7, 14, and 28 were each split (the sampling time corresponded to the different stages of composting, including heating, thermophilic, cooling, and maturity stages), after which they were stored at −80 °C prior to DNA extraction.

### DNA extraction and polymerase chain reaction (PCR) amplification

Total genomic DNA was extracted using an E.Z.N.A^®^ soil DNA kit (Omega Bio-Tek, Norcross, GA, USA) according to the manufacturer’s instructions. The quality and concentration of the purified DNA were determined by Infinite^®^ 200 PRO (TECAN, Sweden). Three copies of each sample were extracted and stored at −20 °C before subsequent analysis.

Real-time qPCR was applied to quantify the presence of 21 ARGs with high abundance that were selected from 274 ARGs. A total of 2 integrase genes, 3 transposable enzyme genes, and 16 s rDNA genes were detected.

Plasmids containing these specific genes were produced by Tianke Biotechnology Co. Ltd (Zhejiang, China) as a standard product with a 10-fold diluted qPCR standard curve, and sterile water used as a negative control. The detected ARGs and MGEs were analyzed further by qPCR using a Bio-Rad IQ5 real-time PCR system (Bio-Rad, USA). The SYBR® Premix Ex Taq™ (TaKaRa, Japan) was used as the fluorescent dye for the qPCR. The qPCR mixture (total volume, 10 µL) contained 5 mL of TB Green Premix Ex Taq II (Tli RNaseH Plus), 0.4 µL of each primer, 0.2 µL of ROX Reference Dye, 1 mL of DNA template, and 3 µL of ddH2O. The thermal cycling steps for qPCR amplification were as follows: an initial denaturation at 95 °C for 10 min, followed by 40 cycles of denaturation at 95 °C for 30 s, annealing for 30 s at the specific temperatures, and then extension at 72 °C for 5 min. Primer specificity was confirmed by melting curves and gel electrophoresis. Each reaction was run in triplicate.

### 16S rRNA gene sequencing

Microbial DNA was extracted from chicken manure samples using an E.Z.N.A.® soil DNA Kit (Omega Bio-tek, Norcross, GA, U.S.) according to manufacturer’s protocols. The final DNA concentration and purification were determined by NanoDrop 2000 UV-vis spectrophotometer (Thermo Scientific, Wilmington, USA), and DNA quality was checked by 1% agarose gel electrophoresis. The V3-V4 hypervariable regions of the bacteria 16S rRNA gene were amplified with primers 338F (5’-ACTCCTACGGGAGGCAGCAG-3’) and 806R (5’-GGACTACHVGGGTWTCTAAT-3’) by thermocycler PCR system (GeneAmp 9700, ABI, USA). The PCR reactions were conducted using the following program: 3 min of denaturation at 95°C, 27 cycles of 30 s at 95°C, 30s for annealing at 55°C, and 45s for elongation at 72°C, and a final extension at 72°C for 10 min. PCR reactions were performed in triplicate 20 μL mixture containing 4 μL of 5 × FastPfu Buffer, 2 μL of 2.5 mM dNTPs, 0.8 μL of each primer (5 μM), 0.4 μL of FastPfu Polymerase and 10 ng of template DNA. The resulted PCR products were extracted from a 2% agarose gel and further purified using an AxyPrep DNA Gel Extraction Kit (Axygen Biosciences, Union City, CA, USA) and quantified using QuantiFluor™-ST (Promega, USA) according to the manufacturer’s protocol.

### Illumina MiSeq sequencing and processing of sequencing data

Purified amplicons were pooled in equimolar and paired-end sequenced (2 × 300) on an Illumina MiSeq platform (Illumina, San Diego, USA) according to the standard protocols by Majorbio Bio-Pharm Technology Co. Ltd. (Shanghai, China).

Raw fastq files were quality-filtered by Trimmomatic and merged by FLASH with the following criteria: (i) reads were truncated at any site receiving an average quality score < 20 over a 50 base pair (bp) sliding window, (ii) sequences whose overlap were longer than 10 bp were merged according to their overlap with mismatch no more than 2 bp, and (iii) sequences of each sample were separated according to barcodes (exactly matching) and primers (allowing 2 nucleotide mismatching), and reads containing ambiguous bases were removed.

Operational taxonomic units (OTUs) were clustered with 97% similarity cutoff using UPARSE (version 7.1 http://drive5.com/uparse/) with a novel “greedy” algorithm that performs chimera filtering and OTU clustering simultaneously. The taxonomy of each 16S rRNA gene sequence was analyzed with a RDP Classifier algorithm (http://rdp.cme.msu.edu/) against the Silva (SSU123) 16S rRNA database using a confidence threshold of 70%.

### Statistical analysis

Analysis of data processing and the generation of plots for the evolution of MGEs used Microsoft Excel 2019. Community barplot, Alpha diversity and community heatmap analysis were performed using the free online platform of Majorbio I-Sanger Cloud Platform (www.i-sanger.com). The variations of persistent ARGs were conducted in R with the package “pheatmap” and Microsoft Excel 2019. The Spearman’s correlation was performed using SPSS 24.0 and *p* <0.05 was considered statistically significant. Network analysis based on the Spearman analysis between ARGs and the bacterial community (based on OTUs) was determined using the Gephi platform (0.9.2).

## Results

### Changes in the bacterial community

The dominant phyla in the chicken manure were distributed among *Firmicutes, Bacteroidetes, Actinobacteria*, and *Proteobacteria*, and they accounted for over 90% of the bacterial population in the chicken manure. After composting, the community structure of bacteria and abundance changed (Fig 1). At the end of the composting period, *Firmicutes* and *Actinobacteria* decreased, and *Bacteroidetes, Deinococcus-Thermus* and *Proteobacteria* increased. At the genus level, we found that there were different compositions among four treatments (Fig 2).

**Fig 1.**
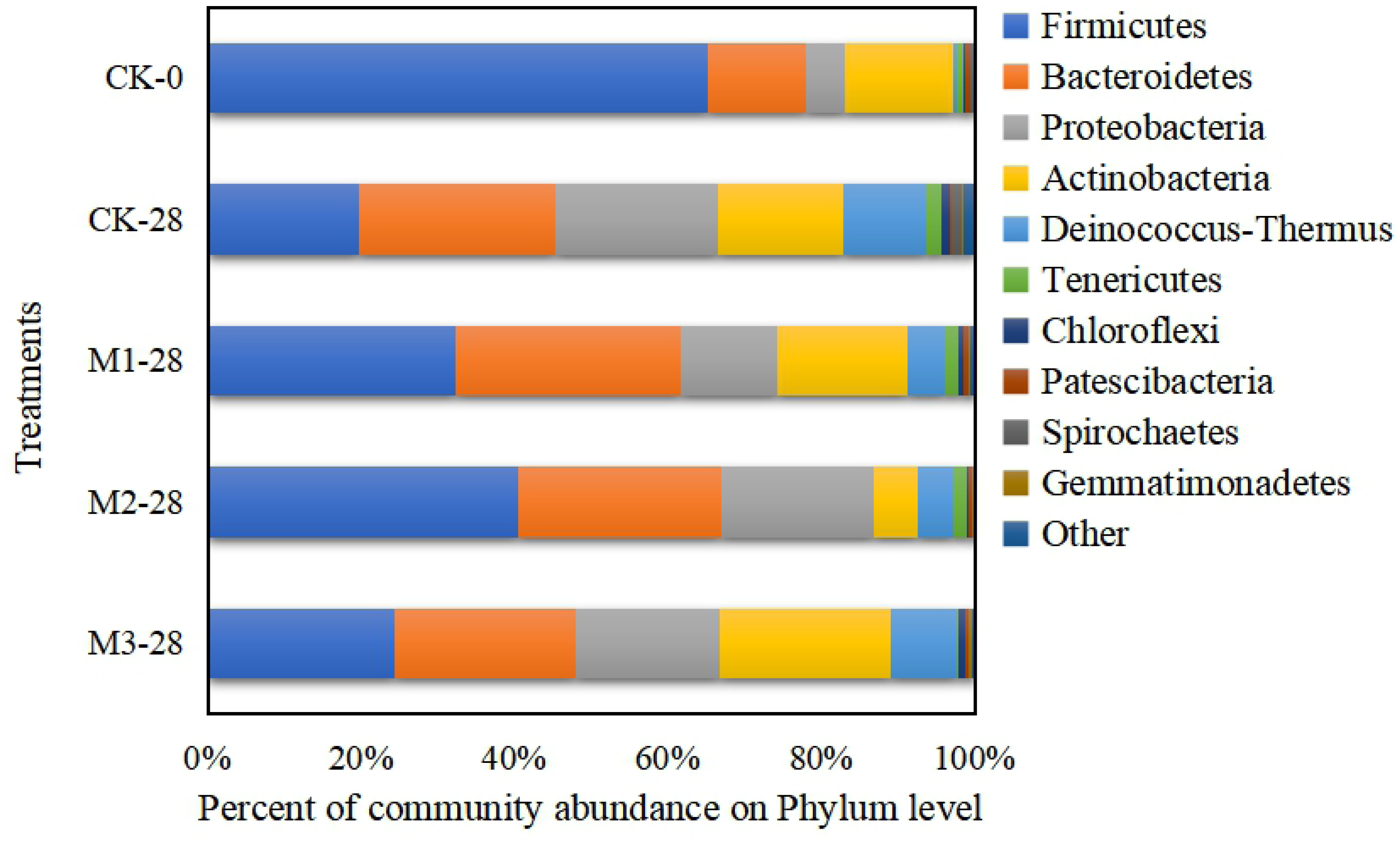
Relative abundance of bacterial phyla of the mine soils in the manure with Chinese herbal medicine residues treatments. The labels under each row denote the treatment name followed by date of compost, for example, CK-28 represents composting samples for the day 28 of control check.

**Fig 2.**
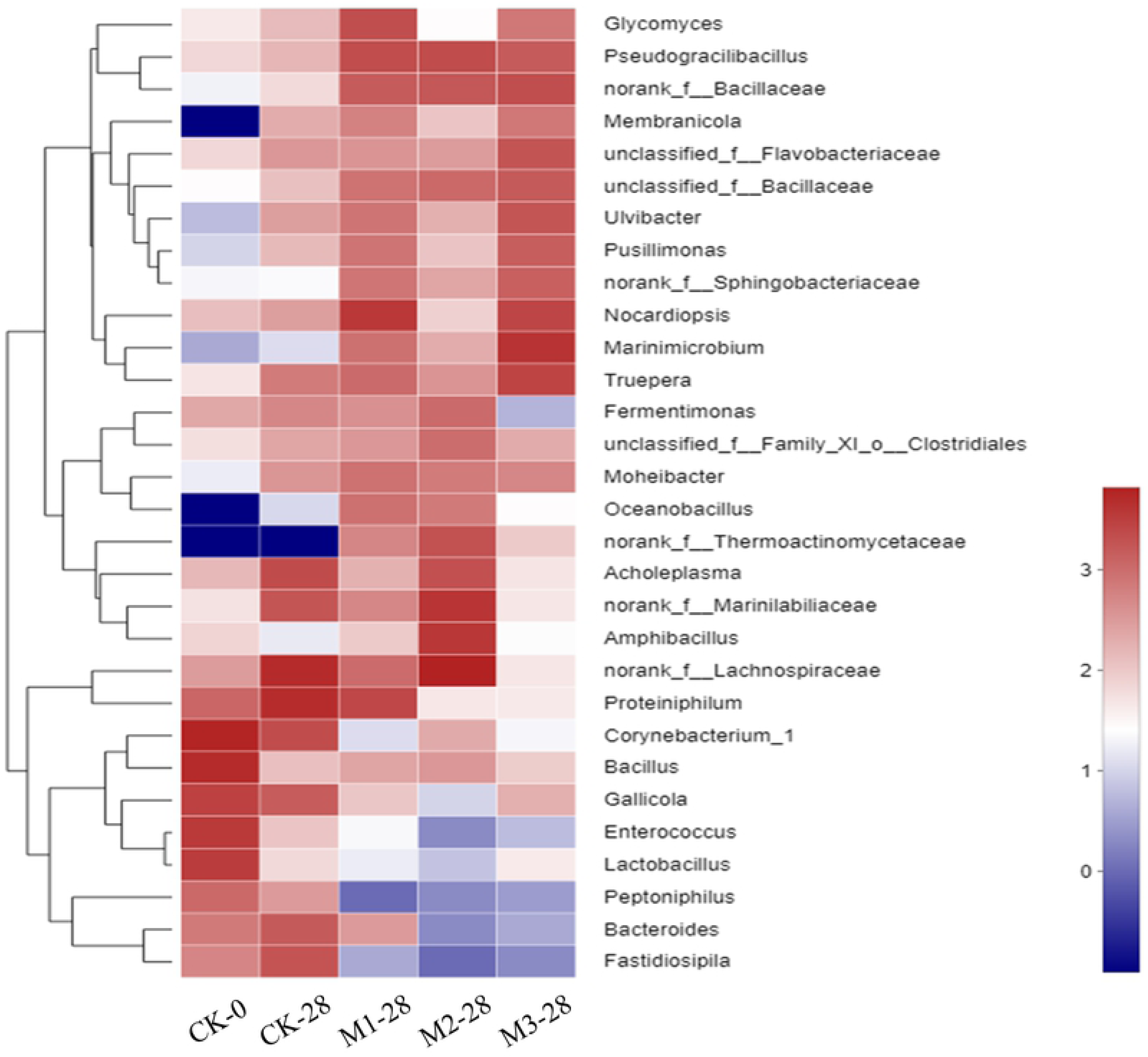
Heatmap showing the relative abundances of the top 30 most abundant genera among all treatments during the composting period. The abscissa is the sample name. The labels under each column denote the treatment name followed by date of compost, for example, CK-3 represents composting samples for the third day of control check. The color blocks in the heatmap represent the relative abundance of a certain item, and the value represented by the color gradient is on the right side of the figure.

Bacterial diversity indices were listed in Table 1. After the end of composting, the Shannon, ACE and Chao1 indices of the treatment (including CK-28) were generally reduced, and the Simpson index increased.

**Table 1.** Alpha diversity index of bacterial community in the manure with Chinese herbal medicine residues treatments.

| Sample | Shannon index | Simpson index | Ace index | Chao index |
| --- | --- | --- | --- | --- |
| CK-0 | 4.89b | 0.02a | 1251.7c | 1287.9d |
| CK-28 | 4.51ab | 0.03a | 987.63b | 1020.42c |
| M1-28 | 4.79b | 0.03a | 916.08ab | 934.77bc |
| M2-28 | 4.08a | 0.06a | 740.21a | 717.44a |
| M3-28 | 3.95a | 0.08a | 795.96ab | 796.4ab |

The different normal letters in the same column indicate significant difference among treatment at 0.05 level. The sample name denotes the treatment name followed by date of compost, for example, CK-28 represents composting samples for the day 28 of control check.

### Variations of the relative abundances of MGEs during composting

The changes of MGEs during the different composting stages are shown in Fig 3. The relative abundances of transposase gene and *int* I 2 decreased with composting, however, the relative abundance of *int* I 1 increased (Fig 3.D). Compared to the control (CK-0), the total relative abundance of MGEs in the treatments with CMHRs.

**Fig 3.**
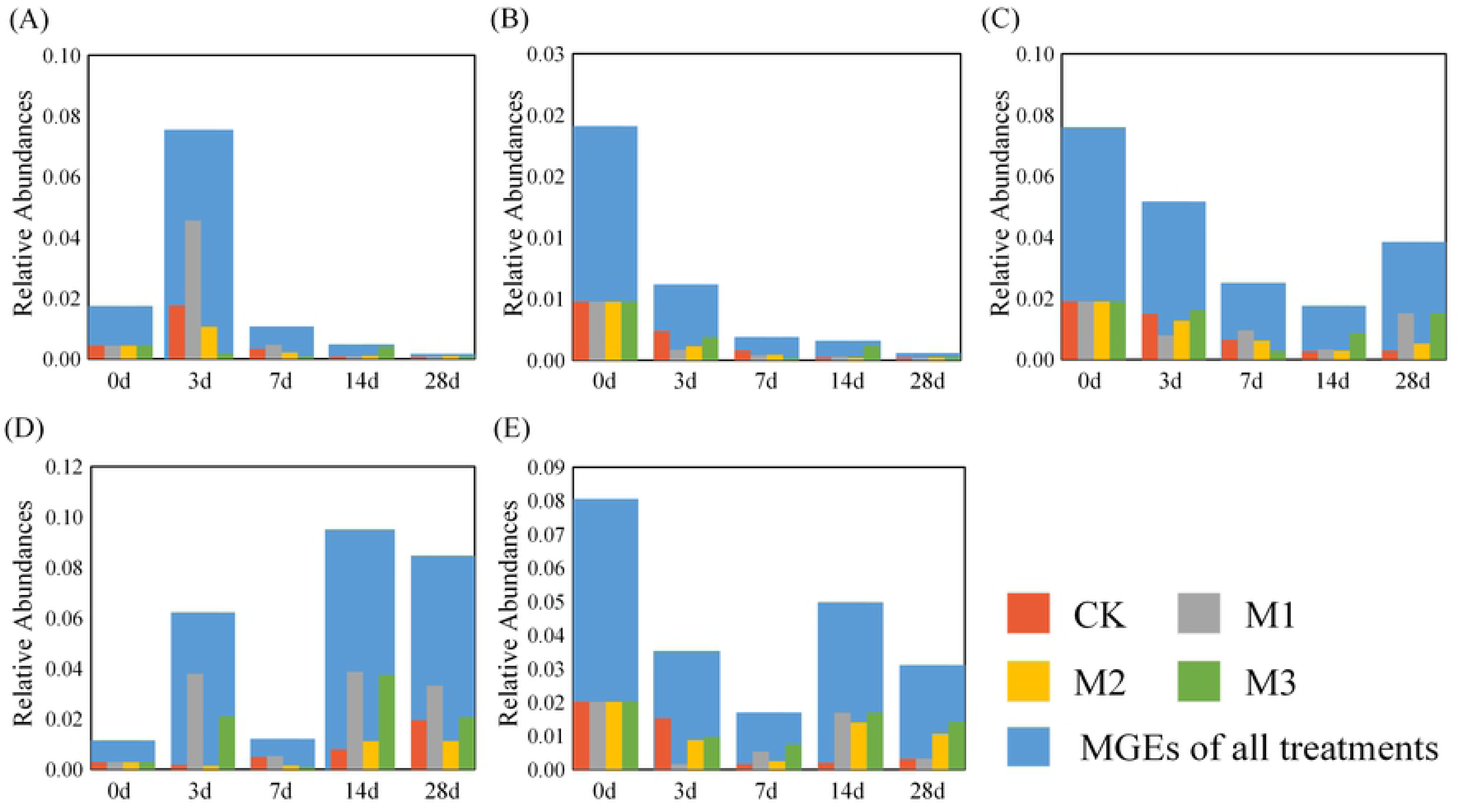
Variations of the relative abundances of *tnp*A (A), *tnp*A-02 (B), *tnp*A-03 (C), *int* I 1 (D), and *int* I 2 (E) during the composting period.

### Variations of the relative abundances of ARGs during composting

ARGs exhibited different fates in different treatments during different composting stages, and the patterns of ARGs during the different composting stages are expressed as heatmaps (Fig 4). Compared with the initial value (0 d), the relative abundance of different ARGs was reduced after 28 days of composting to varying degrees (except sul I). Compared with the control, the relative abundance of ARGs decreased in the three treatments with CMHRs. Especially, the highest removal ratios of ARGs was in M1 with the highest CMHRs (Fig 4B). During composting, the relative abundance of ARGs increased briefly after 3 days of composting (thermophilic stage), and then began to decrease after 14 days (cooling stage).

**Fig 4.**
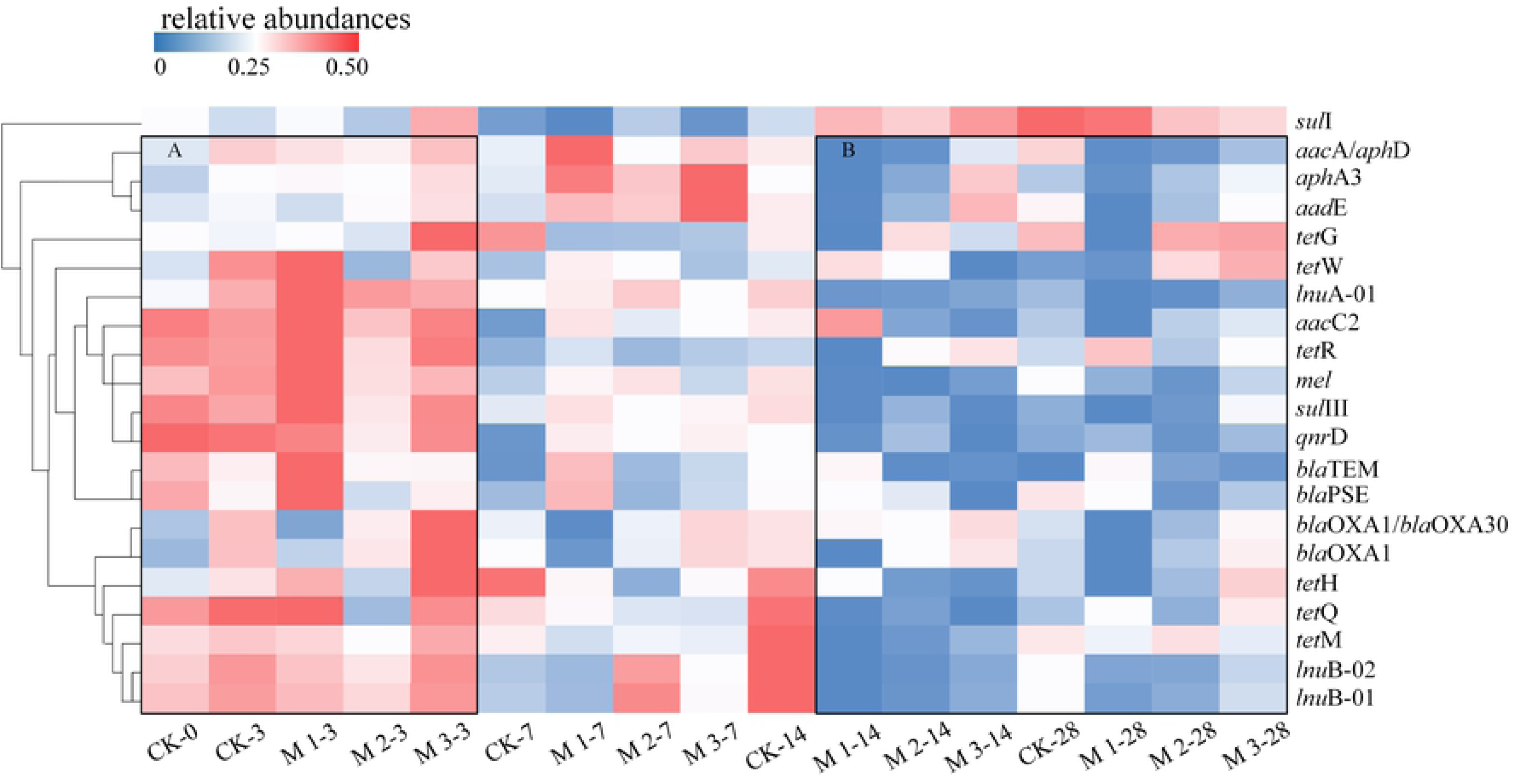
Heatmaps showing the distribution profiles of the relative abundances of antibiotic resistance genes across the different treatments during composting. Black boxes represent different patterns of antibiotic resistance genes: (A) The relative abundance of antibiotic resistance genes increased; (B) The relative abundance of antibiotic resistance genes decreases. The heat scale indicates the relative gene abundance values. The labels under each column denote the treatment name followed by date of compost, for example, CK-3 represents composting samples for the third day of control check.

### Correlation analysis of ARGs and MGEs

Correlations between ARGs and MGEs were analyzed and the results shown in Table 2. For different ARGs, the different correlations with MGEs were demonstrated. *int* I 1 was positively correlated with sulfonamide resistance genes (*p* < 0.01). The transposase gene *tnp*A had a significant positive correlation with the beta lactamase resistance gene and quinolone resistance gene (*p* < 0.05). *tnp*A-02 was positively correlated with the beta lactamase resistance gene and macrolide-linamide (*p* < 0.05). *tnp*A-03 showed no significant correlation with any ARGs.

**Table 2.** Correlations among the abundance of antibiotic resistant genes and mobile gene elements.

|  | *intI1* | *intI2* | *tnpA* | *tnpA*-02 | *tnpA*-03 |
| --- | --- | --- | --- | --- | --- |
| Aminoglycoside-RG | -0.382 | -0.167 | 0.282 | 0.27 | -0.118 |
| Beta Lactamase-RG | -0.137 | 0.127 | 0.507* | 0.539* | 0.277 |
| Macrolide-lincomide-RG | -0.35 | -0.336 | 0.463 | 0.493* | 0.167 |
| Quinolone-RG | -0.453 | -0.017 | 0.569* | 0.466 | 0.426 |
| Sulfonamide-RG | 0.762** | 0.306 | -0.468 | -0.137 | 0.096 |
| Tetracycline-RG | -0.069 | -0.348 | 0.174 | 0.235 | 0.12 |

\*\* Correlation is significant at the 0.01 level (2-tailed).
\* Correlation is significant at the 0.05 level (2-tailed).

### Co-occurrence patterns between ARGs and bacterial community

To explore the co-occurrence relationship between the bacterial community and ARGs, we analyzed the potential host relationships between 50 genera and 6 types ARGs using network analysis. The co-occurrence patterns among ARG subtypes were explored using network inference based on strong (*r* > 0.6) and significant (*p* < 0.01) correlations. Based on the modularity class, the entire network could be parsed into 7 major modules, with 60 of 70 total vertices occupied by the 3 largest modules: Modules I, II, and III (Fig 5A). Network analysis obtained 70 nodes and 302 edges, with 20 ARGs and 50 genera of bacteria distributed in 9 phyla, mainly Firmicutes (Fig 5B). Each bacterial taxon occurred simultaneously with more than one ARGs and each ARG also had more than one potential host bacteria. *Enterococcus* may be the host bacterium of *aph*A3, *blo*OXA1/*bla*OXA30, and *tet*H. *tet*R and *qnr*D only had one potential host bacterial taxon, *Fermentimonas* and *Achleplasma*, respectively. More than one genus of bacteria was the host of the same ARGs, for example, *sul* I was found in *Bacteroides, Fastidiospila, Pseudomonas*, and *Proteiniphilum*.

**Fig 5.**
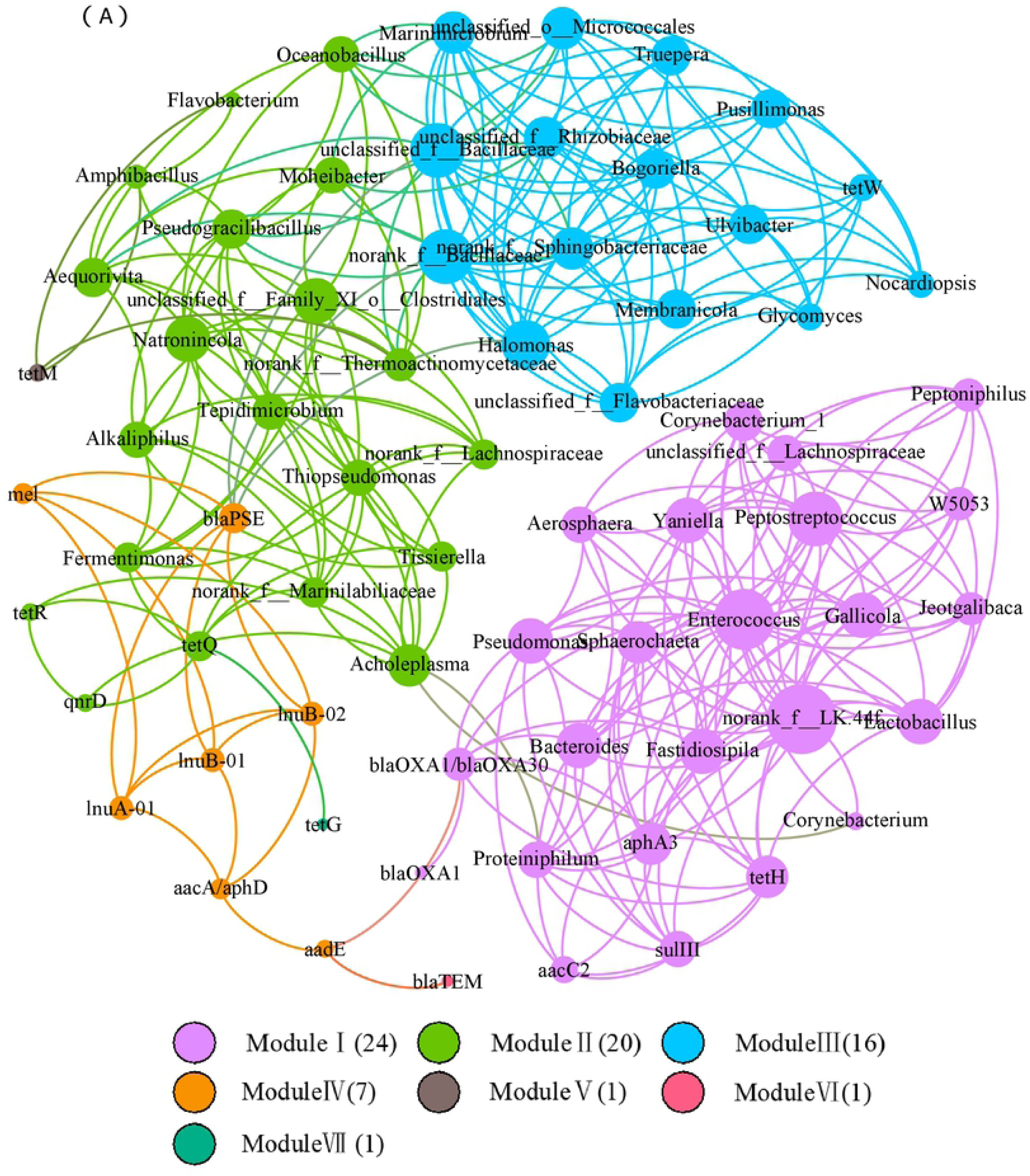
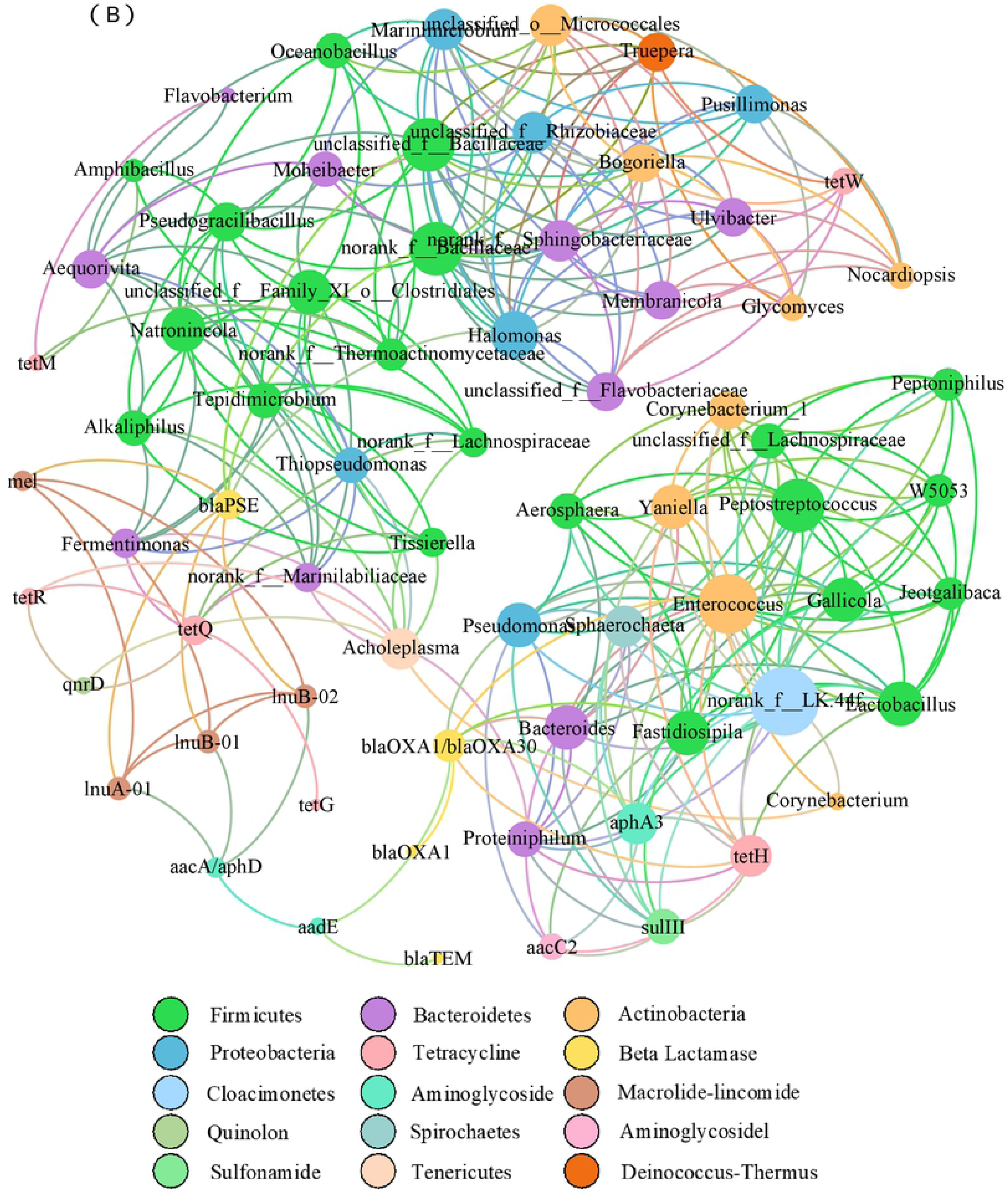
Network analysis based on the co-occurrence of antibiotic resistance genes and their potential host bacteria. The nodes are colored according to antibiotic resistance genes types and genus (A). The nodes are colored according to modularity class (B). The size of each node is proportional to the number of connections, that is, the degree.

## Discussion

Microorganisms are carriers of ARGs, so a change of bacterial community will lead to a change of abundance of ARGs and affect their migration in the environment. We detected the dominant bacteria in chicken manure as being *Firmicutes, Bacteroidetes, Actinobacteria*, and *Proteobacteria* (Fig 1), which is similar to that found by previous studies (Huang et al., 2018; Qian et al., 2016;Zhang et al., 2018). Previous studies confirmed that *Firmicutes* may be the host bacteria for carrying and spreading ARGs (Huerta et al., 2013), furthermore, the fate of ARGs has been significantly correlated with microbial community succession changes, especially the reduction in *Clostridiales* (belongs to the phylum *Firmicutes*) (Wang et al., 2017). *Firmicutes* in our study decreased substantially after composting which could partly explain the decrease of the relative abundance of ARGs after the composting. Such result was also supported by which the domain host bacteria carrying ARGs was Firmicutes through the Network analysis (Fig 5B). *Bacteroidetes* and *Proteobacteria* increased after composting (Fig 1), which is coherent to previous studies of sludge composting system (Huang et al., 2018). The bacterial diversity and abundance decreased after composting, but the difference in whether or not to add CMHRs was not significant (Table 1), which would due to the high temperatures during the composting period reducing the number of drug-resistant bacteria entering the environment.

The abundance of ARGs and MGEs decreased after composting (except for *sul* I and *int* I 1) (Figs 2 and 3), which suggested that the horizontal gene transfer was inhibited and ARGs transmission was reduced by decreasing the abundance of MGEs. The removal ratio of ARGs was high in M1 with the highest CMHRs (Fig 4). *sul*I kept high abundance after composting which could due to the thermal stability of sulfonamide. Such result is consistent with previous studies (Su et al., 2015; Lin et al., 2016; Qian et al., 2016). The variation of ARGs during composting were related to different mechanisms and environmental pressures (Wright et al., 2007; Li et al., 2017) which could explain the different trends for different types of ARGs during composting.

The transposase gene and *int*I2 effectively decreased after composting, but *int*I1 increased significantly in this study. The increase on *int*I1 and the decrease on *int*I2 during the later composting stage were also reported by Zhang et al. (2017). The decrease of *int*I2 was due to the sensitivity of integrase genes to high temperature (Ghosh et al., 2009; Ma et al., 2011; Selvam et al., 2012). Correlation analysis (Table 2) showed that *int* I 1 was positively correlated with sulfonamide resistance genes, which can be explained by the fact that *sul*1 genes are usually located on class 1 integrons and are associated with other resistance genes (Antunes et al., 2005). In our study, transposons were significantly positively correlated with the beta lactamase and quinolone resistance genes. Previous studies showed that *tnp*A is significantly correlated with the tetracycline resistance gene (Zhu et al., 2013) and aminoglycoside resistance gene (Chen et al., 2016).

## Conclusion

Adding CMHRs as compost-bulking agents, the majority of ARGs and MGEs were removed (except for *sul* I and *int* I 1) after composting, and the removal rate was in the order of M1 > M2 > M3 > CK. *sul* I and *int* I 1 showed a significant positive correlation (*p* < 0.01). After composting, the microbial community structure of the different treatments changed, however, similar trends were observed in the bacterial community structure among the different CMHR treatment. Network analysis and heatmap analysis showed that the increase and decrease of bacteria were similar to that of ARGs, indicating that they might be the host bacteria of ARGs and thus affect the removal of ARGs.

## Acknowledgement

This work was financially supported by the Natural Science Foundation of China (41271530); the innovation projects of Shanxi Education Department of China (2018); Shanxi Key R&D Program (Social Development Field) Project (201803D31024); and Shanxi Higher Education Institution Project: Ecological Remediation of Soil Pollution Disciplines Group (20181401).

## References

Antunes P, Machado J, Sousa JC, Peixe L. Dissemination of sulfonamide resistance genes (sul1, sul2, and sul3) in portuguese salmonella enterica strains and relation with integrons. Antimicrob Agents Ch. 2005; 49:836–839. https://doi.org/10.1128/AAC.49.2.836-839.2005

Chang YJ, Tang MH, Cheng WD. Experimental study on effects of chinese medicine residues on soil amelioration. Modern Agricultural Science and Technology. 2010; 5:255–256. https://doi.org/10.3969/j.issn.1007-5739.2010.05.182

Chen J, Yu Z, Michel FC, Jr., Wittum T, Morrison M. Development and application of real-time pcr assays for quantification of erm genes conferring resistance to macrolides-lincosamides-streptogramin b in livestock manure and manure management systems. Appl Environ Microbiol. 2007; 73:4407–4416. https://doi.org/10.1128/AEM.02799-06

Chen Q, An X, Li H, Su J, Ma Y, Zhu YG. Long-term field application of sewage sludge increases the abundance of antibiotic resistance genes in soil. Environ Int. 2016; 92-93:1–10. https://doi.org/10.1016/j.envint.2016.03.026

Cui E, Wu Y, Zuo Y, Chen H. Effect of different biochars on antibiotic resistance genes and bacterial community during chicken manure composting. Bioresour Technol. 2016; 203:11–17. https://doi.org/10.1016/j.biortech.2015.12.030

Duan M, Gu J, Wang X, Li Y, Zhang S, Yin Y, Zhang R. Effects of genetically modified cotton stalks on antibiotic resistance genes, inti1, and inti2 during pig manure composting. Ecotoxicol Environ Saf. 2018; 147:637–642. https://doi.org/10.1016/j.ecoenv.2017.09.023

Ghosh S, Ramsden SJ, LaPara TM. The role of anaerobic digestion in controlling the release of tetracycline resistance genes and class 1 integrons from municipal wastewater treatment plants. Appl Microbiol Biotechnol. 2009; 84:791–796. https://doi.org/10.1007/s00253-009-2125-2

Han XM, Hu HW, Chen QL, Yang LY, Li HL, Zhu YG, Li XZ, Ma YB. Antibiotic resistance genes and associated bacterial communities in agricultural soils amended with different sources of animal manures. Soil Biol Biochem. 2018; 126:91–102. https://doi.org/10.1016/j.soilbio.2018.08.018

Huang K, Xia H, Wu Y, Chen J, Cui G, Li F, Chen Y, Wu N. Effects of earthworms on the fate of tetracycline and fluoroquinolone resistance genes of sewage sludge during vermicomposting. Bioresour Technol. 2018; 259:32–39. https://doi.org/10.1016/j.biortech.2018.03.021

Huerta B, Marti E, Gros M, Lopez P, Pompeo M, Armengol J, Barcelo D, Balcazar JL, Rodriguez-Mozaz S, Marce R. Exploring the links between antibiotic occurrence, antibiotic resistance, and bacterial communities in water supply reservoirs. Sci Total Environ. 2013; 456:161–170. https://doi.org/10.1016/j.scitotenv.2013.03.071

Laguë C, Landry H, Roberge M. Engineering of land application systems for livestock manure: A review. Canadian Biosystems Engineering. 2005; 47:6.17-6.28. https://doi.org/10.1016/j.agee.2013.08.009

Li H, Duan M, Gu J, Zhang Y, Qian X, Ma J, Zhang R, Wang X. Effects of bamboo charcoal on antibiotic resistance genes during chicken manure composting. Ecotoxicol Environ Saf. 2017; 140:1–6. https://doi.org/10.1016/j.ecoenv.2017.01.007

Li JJ, Xin ZH, Zhang YZ, Chen JW, Yan JX, Li HJ, Hu HW. Long-term manure application increased the levels of antibiotics and antibiotic resistance genes in a greenhouse soil. Appli Soil Ecol. 2017; 121:193–200. https://doi.org/10.1016/j.apsoil.2017.10.007

Lin H, Wang J, Sun W, Fu J, Chen H, Ma J. Interaction between sulfonamide antibiotics fates and chicken manure composting. Huan jing ke xue= Huanjing kexue.2016; 37:1993–2002. https://doi.org/10.13227/j.hjkx.2016.05.050

Ma YJ, Wilson CA, Novak JT, Riffat R, Aynur S, Murthy S, Prudens A. Effect of various sludge digestion conditions on sulfonamide, macrolide, and tetracycline resistance genes and class i integrons. Environ Sci Technol.2011; 45:7855–7861. https://doi.org/10.1021/es200827t

Martinez JL, Sanchez MB, Martinez-Solano L, Hernandez A, Garmendia L, Fajardo A, Alvarez-Ortega C. Functional role of bacterial multidrug efflux pumps in microbial natural ecosystems. FEMS Microbiol Rev. 2009; 33:430–449. https://doi.org/10.1111/j.1574-6976.2008.00157.x

Qian X, Sun W, Gu J, Wang XJ, Sun JJ, Yin YN, Duan ML. Variable effects of oxytetracycline on antibiotic resistance gene abundance and the bacterial community during aerobic composting of cow manure. J Hazard Mater. 2016; 315:61–69. https://doi.org/10.1016/j.jhazmat.2016.05.002

Qiu HG, Liao SP, Jing Y, Luan J. [regional differences and development tendency of livestock manure pollution in china]. Huan Jing Ke Xue= Huanjing kexue. 2013; 34:2766–2774. https://doi.org/10.13227/j.hjkx.2013.07.010

Selvam A, Xu D, Zhao Z, Wong JW. Fate of tetracycline, sulfonamide and fluoroquinolone resistance genes and the changes in bacterial diversity during composting of swine manure. Bioresour Technol. 2012; 126:383–390. https://doi.org/10.1016/j.biortech.2012.03.045

Su JQ, Wei B, Ou-Yang WY, Huang FY, Zhao Y, Xu HJ, Zhu YG. Antibiotic resistome and its association with bacterial communities during sewage sludge composting. Environ Sci Technol. 2015; 49:7356–7363. https://doi.org/10.1021/acs.est.5b01012

Tien YC, Li B, Zhang T, Scott A, Murray R, Sabourin L, Marti R, Topp E. Impact of dairy manure pre-application treatment on manure composition, soil dynamics of antibiotic resistance genes, and abundance of antibiotic-resistance genes on vegetables at harvest. Sci Total Environ. 2017; 581:32–39. https://doi.org/10.1016/jscitotenv.2016.12.138

Wang H, Sangwan N, Li HY, Su JQ, Oyang WY, Zhang ZJ, Gilbert JA, Zhu YG, Ping F, Zhang HL. The antibiotic resistome of swine manure is significantly altered by association with the musca domestica larvae gut microbiome. Isme J. 2017; 11:100–111. https://doi.org/10.1038/ismej.2016.103

Wang YQ, Wu XQ, Zhu TT, Ma QG, Chen HG. Study on utilization of solid slag compost of Chinese medicinal herbal. Journal of Chinese Medicinal Materials. 2008; 31:1622–1624. https://doi.org/10.13863/j.issn1001-4454.2008.11.005

Wright GD. The antibiotic resistome: The nexus of chemical and genetic diversity. Nat Rev Microbiol. 2007; 5:175–186. https://doi.org/10.1038/nrmicro1614

Zhang J, Chen M, Sui Q, Tong J, Jiang C, Lu X, Zhang Y, Wei Y. Impacts of addition of natural zeolite or a nitrification inhibitor on antibiotic resistance genes during sludge composting. Water Res. 2016; 91:339–349. https://doi.org/10.1016/j.watres.2016.01.010

Zhang J, Lin H, Ma J, Sun W, Yang Y, Zhang X. Compost-bulking agents reduce the reservoir of antibiotics and antibiotic resistance genes in manures by modifying bacterial microbiota. Sci Total Environ. 2019; 649:396–404. https://doi.org/10.1016/j.scitotenv.2018.08.212

Zhang L, Gu J, Wang X, Sun W, Yin Y, Sun Y, Guo A, Tuo X. Behavior of antibiotic resistance genes during co-composting of swine manure with chinese medicinal herbal residues. Bioresour Technol. 2017; 244:252–260. https://doi.org/10.1016/j.biortech.2017.07.035

Zhang QQ, Ying GG, Pan CG, Liu YS, Zhao JL. Comprehensive evaluation of antibiotics emission and fate in the river basins of china: Source analysis, multimedia modeling, and linkage to bacterial resistance. Environ Sci Technol. 2015; 49:6772–6782. https://doi.org/10.1021/acs.est.5b00729

Zhang R, Gu J, Wang X, Li Y, Zhang K, Yin Y, Zhang X. Contributions of the microbial community and environmental variables to antibiotic resistance genes during co-composting with swine manure and cotton stalks. J Hazard Mater. 2018; 358:82–91. https://doi.org/10.1016/j.jhazmat.2018.06.052

Zhang Y, Li H, Gu J, Qian X, Yin Y, Li Y, Zhang R, Wang X. Effects of adding different surfactants on antibiotic resistance genes and inti1 during chicken manure composting. Bioresour Technol. 2016; 219:545–551. https://doi.org/10.1016/j.biortech.2016.06.117

Zhao W, Wang B, Yu G. Antibiotic resistance genes in china: Occurrence, risk, and correlation among different parameters. Environ Sci Pollut R. 2018; 25:21467–21482. https://doi.org/10.1007/s11356-018-2507-z

Zhou QX, Luo Y, Wang ME. Environmental residues and ecotoxicity of antibiotics and their resistance gene pollution: A review. Asian Journal of Ecotoxicology. 2007; 2:243–251. https://doi.org/10.1007/s11767-005-0212-9

Zhu YG, Johnson TA, Su JQ, Qiao M, Guo GX, Stedtfeld RD, Hashsham SA, Tiedje JM. Diverse and abundant antibiotic resistance genes in chinese swine farms. Proc Natl Acad Sci USA. 2013; 110:3435–3440. https://doi.org/10.1073/pnas.1222743110

